# Upper Limit Efficiency Estimates for Electromicrobial Production of Drop-In Jet Fuels

**DOI:** 10.1101/2022.10.12.511952

**Authors:** Timothy J. Sheppard, David Specht, Buz Barstow

## Abstract

Microbes which participate in extracellular electron uptake or H_2_ oxidation have an extraordinary ability to manufacture organic compounds using electricity as the primary source of metabolic energy. So-called electromicrobial production could be of particular value in the efficient production of hydrocarbon blends for use in aviation. Because of exacting standards for fuel energy density and the costs of new aviation infrastructure, liquid hydrocarbon fuels will be necessary for the foreseeable future, precluding direct electrification. Production of hydrocarbons using electrically-powered microbes employing fatty acid synthesis-based production of alkanes could be an efficient means to produce drop-in replacement jet fuels using renewable energy. Here, we calculate the upper limit electrical-to-energy conversion efficiency for a model jet fuel blend containing 85% straight-chain alkanes and 15% terpenoids. When using the Calvin cycle for carbon-fixation, the energy conversion efficiency is 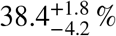 when using extracellular electron uptake for electron delivery and 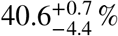 when using H_2_-oxidation. The efficiency of production of the jet fuel blend can be raised to 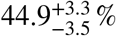 when using the Formolase formate-assimilation pathway and H_2_-oxidation, and to 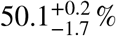 with the Wood-Ljungdahl pathway. The production efficiency can be further raised by swapping the well-known ADO pathway for alkane termination with for the recently discovered MCH pathway. If these systems were were supplied with electricity with a maximally-efficient silicon solar photovoltaic, even the least efficient would exceed the maximum efficiency of all known forms of photosynthesis.

## Introduction

While global aviation only represents 2.4% of anthropogenic CO_2_ emissions^1^, decarbonization of this industry is especially challenging, both because of the high capital costs of new aircraft as well as the requirement for highly energy dense fuels. As a result, the aviation industry will continue to be dependent on liquid fuels for decades to come despite advances in the electrification of other forms of transportation^2^. Ambitious decarbonization goals, for example those set by the International Air Transport Association for net-zero carbon emissions by 2050^3^, rely predominantly on the use of biologically-derived sustainable aviation fuels (SAF). However, it will be challenging to achieve even 2% usage of SAF by 2025^2^, and dramatic advancements in development of these fuels is essential in order to combat anthropogenic climate change^4^. Uncertainty in regulation of carbon emissions, especially in the United States^5^, as well as volatility in jet fuel prices^6^, will continue to drive unpredictability in the future of SAF biofuels. However, we believe that the fundamental issue – the requirement for substantial agricultural inputs – can be addressed through the use of electromicrobial production (EMP), in which electrically-derived inputs are used as a feedstock to drive microbial output^7−11^, in an overall process we term EMP-to-Jet.

Biofuels have long been proposed as a ‘drop-in’ solution to volatility in petroleum-based fuel prices and as a means to reduce GHG emissions. So-called ‘1^st^ generation’ fuels, such as those in the US derived from corn, compete directly with agriculture, have dubious GHG reduction benefits, and do not represent a sustainable path forward due to pressures on arable land usage^12^. Despite advances in ‘2^nd^ generation’ biofuels – derived from non-food oil crops and waste oils – and ‘3^rd^ generation’ biofuels – derived from photosynthetic algae – neither has yet proven to be commercially mainstream, although numerous facilities using diverse low-value feedstocks are anticipated to go online in 2023 and 2024^2^. Further, while first and second-generation biofuels do generally have lower GHG emissions than the use of traditional petroleum-based fuels, this is predicated on the absence of land-use changes^13^, which would seem to be unlikely to be achievable in practice. Full life cycle analyses of current 3^rd^ generation technologies show that they emit more GHG than even petroleum fuels^13^. Achieving emissions targets while simultaneously minimizing land use changes will require substantial advancements, and novel biotechnological solutions provided by synthetic biology may be an answer.

Here, we propose using native mechanisms for hydrogen oxidation^7,8^ or extracellular electron uptake (EEU)^14^ as the sole source of metabolic energy used to drive jet fuel hydrocarbon production (**Figs. 1A** to **C**). Electrical flow provides the reducing power required for metabolism, and carbon fixation is performed completely enzymatically (*e*.*g*., by the Calvin-Bassham-Benson (CBB or Calvin) cycle) or through the assimilation of electrochemically-synthesized C_1_ compounds, such as formate^15,16^ (*e*.*g*., by the Formolase pathway^17^). Theoretical analysis demonstrates that EMP is dramatically more efficient than the use of photosynthesis, both thermodynamically as well as in terms of land use^11,18−21^. Further, while photosynthesizing cyanobacteria are promising in principle, they remain very difficult to engineer^22^. By contrast, at least one microbe capable of EEU can be just as engineerable as molecular biology workhorses: the fast-growing microbe *Vibrio natriegens*, recently demonstrated to be capable of EEU^23^, is often considered a next-generation replacement for *E. coli* due to its ease of engineering^24,25^.

**Figure 1.**
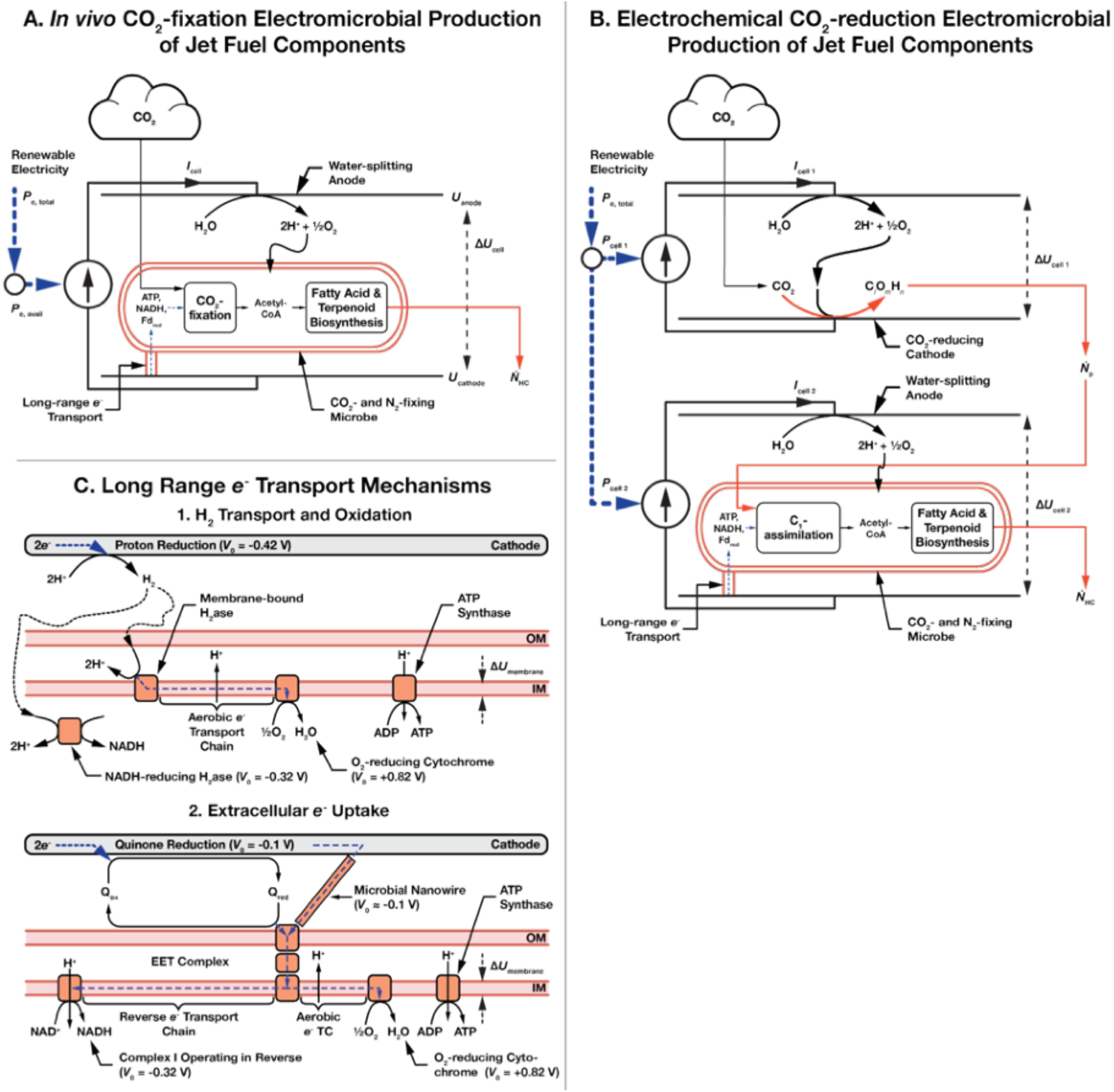
Schematic of electromicrobial jet fuel production systems. Carbon fixation for electromicrobial production could take the form of a fully *in vivo* system (**A**), in which the microbe both fixes carbon and produces the hydrocarbon product, or (**B**), in which an abiotic cell is first used to fix carbon into a lightly-reduced compound such as formate prior to use in a second bio-electrochemical cell used for hydrocarbon production. (**C**) Mechanisms by which electricity sources can be used to power microbial production, using either H_2_ oxidation or extracellular electron uptake. In the first, H_2_ is electrochemically reduced on a cathode, transferred to the microbe by diffusion or stirring, and is enzymatically oxidized. In the second mechanism, extracellular electron uptake (EEU), *e*-are transferred along a microbial nanowire (part of a conductive biofilm), or by a reduced medium potential redox shuttle like a quinone or flavin, and are then oxidized at the cell surface by the extracellular electron transfer (EET) complex. From the thermodynamic perspective considered in this article, these mechanisms are equivalent. Electrons are then transported to the inner membrane where reverse electron transport is used to regenerate NAD(P)H, reduced Ferredoxin (not shown), and ATP^14,34^.

Ethanol based biofuels are often considered for ground transportation but are unsuitable for use in aviation due to insufficient energy density. There are two major categories of compounds which could be manufactured using EMP as components of a drop-in jet fuel substitute: fatty acid-derived alkanes^26^ (**Fig. 2**), and terpenoids^27^ (**Fig. 3**). Here, we will focus on engineering pathways for production of both, with an eye towards producing a blended fuel.

**Figure 2.**
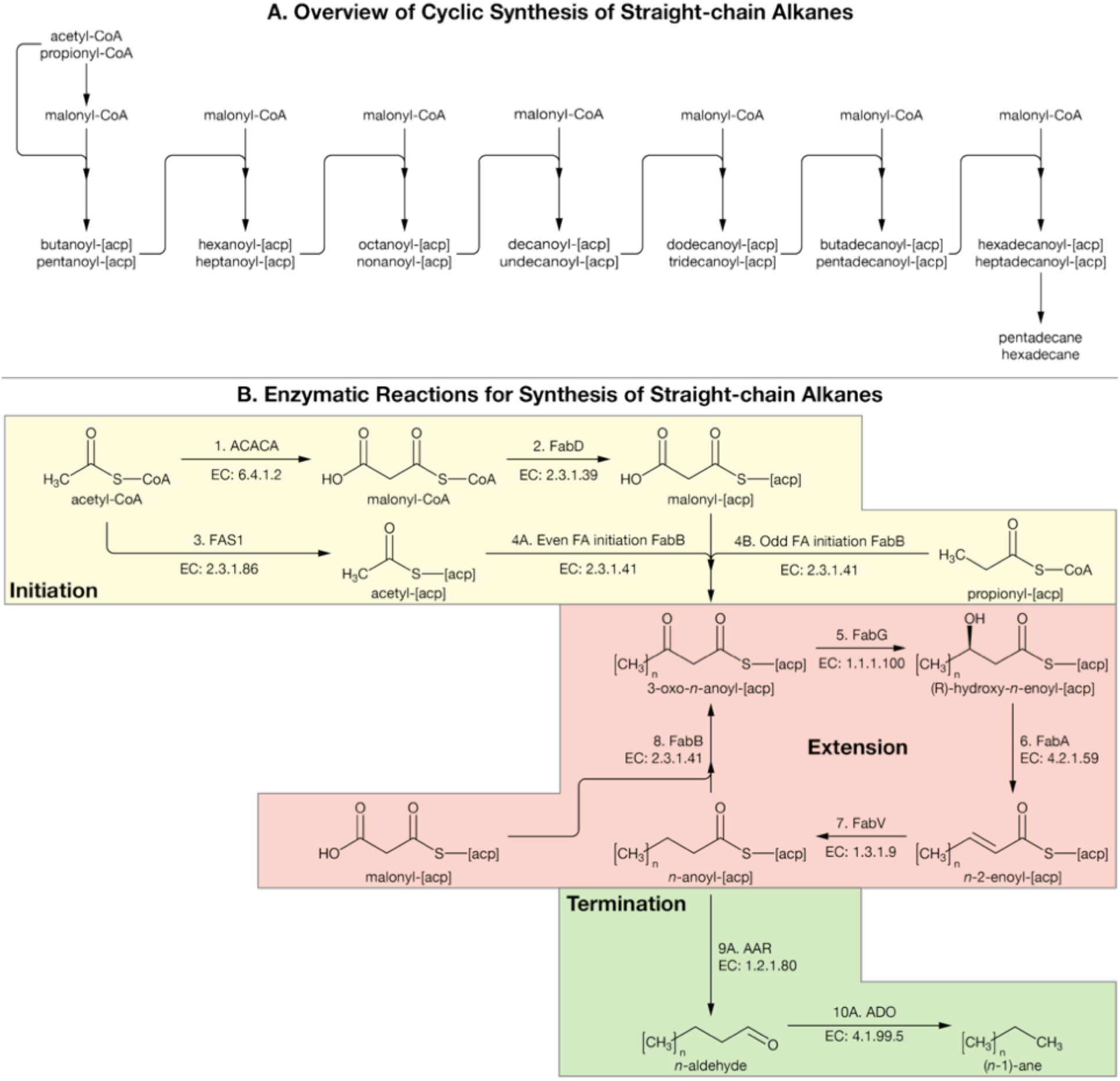
Synthesis pathways for straight-chain alkane components of jet fuel blend. (**A**) *In vivo* production is accomplished via cyclic elongation of fatty acids prior to termination to produce an alkane of the desired length. Note, if the cycle is initiated with propionyl-CoA and malonyl-CoA the fatty acid synthesis cycle will produce odd chain length fatty acids (leading to even alkanes). On the other hand, if the cycle is initiated with acetyl-CoA and malonyl-CoA it will produce even numbered fatty acids (leading to odd alkanes). (**B**) Details on enzymatic reactions for Type II Fatty Acid Synthesis (FAS). Full details on the generalized (for any odd or even chain length) enzymatic reactions can be found in **Table 2**. In this figure we only show alkane termination by the well-studied ADO (Aldehyde Deformolating Oxygenase) pathway. Reactions for the less well-known FAP (Fatty Acid Photodecarboxylase) pathway are shown in **Table 2**. ACACA: Acetyl-CoA Carbonic Anhydrase; FA: Fatty Acid; AAR: Aceto-Aldehyde Reductase. Reactions for Type II Fatty Acid Synthesis are taken from the Kyoto Encyclopedia of Genes and Genomes (KEGG)^45-47^. Reactions for conversion of acetyl-CoA to propionyl-CoA are shown in **Table S4**.

**Figure 3.**
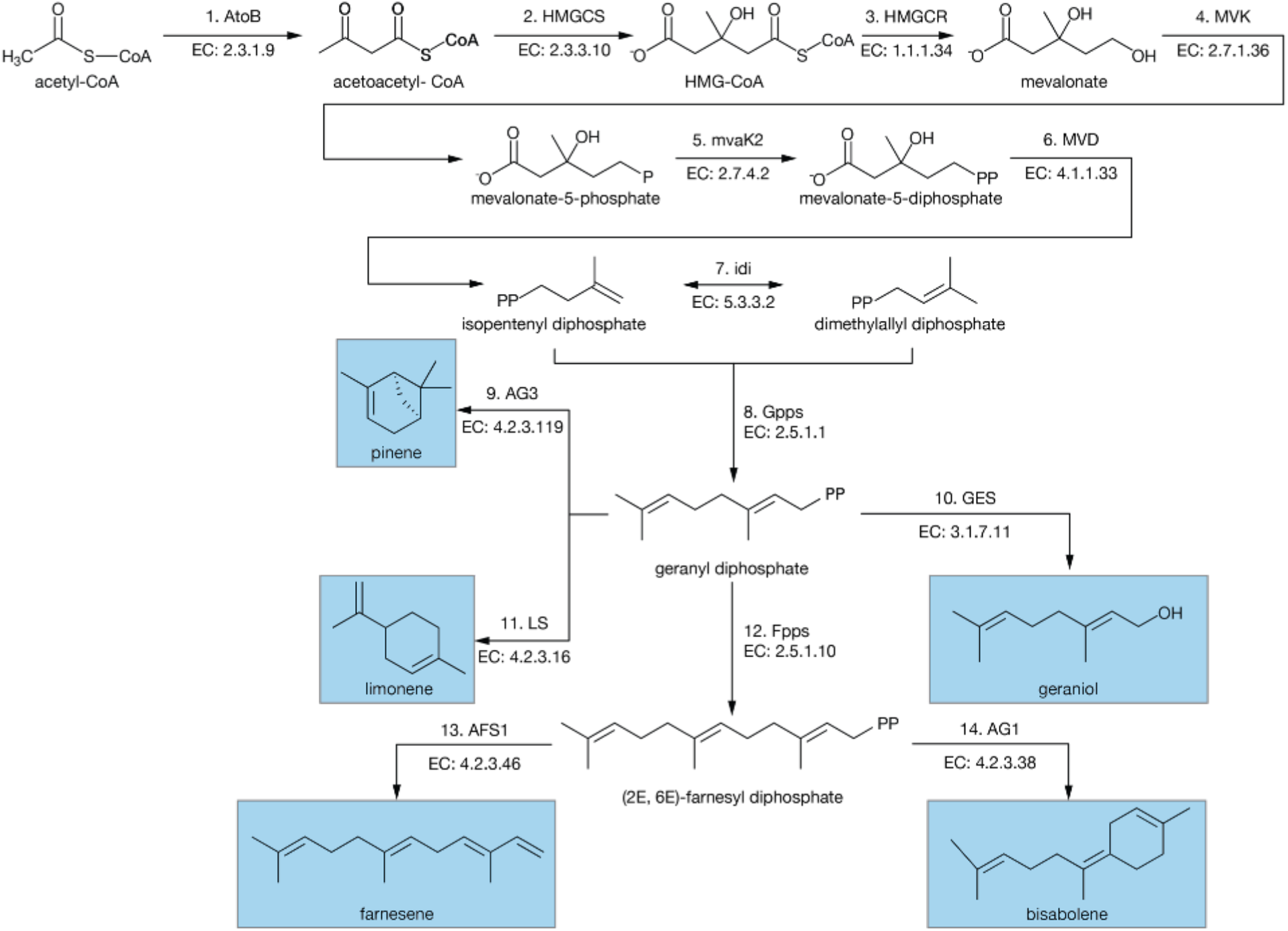
Synthesis pathways for aromatic components of jet fuel blend. Adapted from Adesina *et al*. ^48^. using information from Cheon *et al*.^49^ with additional pathways from Peralta-Yahya *et al*.^50^. Numbers for reactions correspond to entries in **Table 1**.

Alkanes offer desirable properties when burned and are the dominant component in most existing jet fuels. However, a pure aliphatic, alkane fuel will not satisfy current aviation fuel standards. Aviation fuels are not a homogeneous mixture and are standardized not by their chemical composition, but rather by their physical properties. Globally, the most commonly used jet fuel is Jet A-1, with standards set by ASTM (American Society for Testing and Materials) International via standard D1655^28^. These standards balance fuel attributes which are essential for safe aircraft operation. For example, drop-in jet fuels which are purely aliphatic may cause shrinkage of nitrile O-rings used in fuel management^29^, and as a result there is a minimum 8% aromatic content. Conversely, aliphatic fuels burn cleaner, as the addition of aromatics increases soot formation, so there is a 25% aromatic upper limit. Perhaps the most constrictive requirement of ASTM D1655 is the density (775.0 to 840.0 kg m^−3^), which again effectively prohibits a pure alkane fuel^30^.

**Table 1.** Electromicrobial jet fuel production model parameters. Model parameters used in this article are based upon model parameters used in a previous analysis of the electromicrobial production of the biofuel butanol^11^. A sensitivity analysis was performed for all key parameters in this work^11^.

| Parameter | Symbol | 1. H<sub>2</sub> | 2. EEU | 3. H<sub>2</sub> with Formate | 4. EEU with Formate |
| --- | --- | --- | --- | --- | --- |
| **Electrochemical Cell Parameters** |  |  |  |  |  |
| Input solar power (W) | $P_{\gamma}$ | 1,000 | 1,000 | 1,000 | 1,000 |
| Total available electrical power (W) | $P_{e, \text{ total}}$ | 330 | 330 | 330 | 330 |
| CO<sub>2</sub>-fixation method |  | Enzymatic |  | Electrochemical |  |
| Electrode to microbe mediator |  | H<sub>2</sub> | EEU | H<sub>2</sub> | EEU |
| Cell 1 cathode std. potential (V) | $U_{\text{cell 1, cathode, 0}}$ | N/A | | 0.82 [Torella2015a] | |
| Cell 1 cathode bias voltage (V) | $U_{\text{cell 1, cathode, bias}}$ | N/A | | 0.47 [Liu2016a] | |
| Cell 1 anode std. potential (V) | $U_{\text{cell 1, anode, 0}}$ | N/A | | -0.43 [Yishai2017a, Zhang2018a] | |
| Cell 1 anode bias voltage (V) | $U_{\text{cell 1, anode, bias}}$ | N/A | | 1.3 [White2014a] | |
| Cell 1 voltage (V) | $\Delta U_{\text{cell 1}}$ | N/A | | 3.02 | |
| Cell 1 Faradaic efficiency | $\zeta_{I1}$ | N/A | | 0.8 [Rasul2019a] | |
| Carbons per primary fixation product | $v_{Cr}$ | N/A | | 1 | |
| $e^-$ per primary fixation product | $v_{er}$ | N/A | | 2 | |
| Cell 2 (Bio-cell) anode std. potential (V) | $U_{\text{cell 2, anode, 0}}$ | -0.41 [Torella2015a] | -0.1 [Bird2011a, Firer-Sherwood2008a] | -0.41 | -0.1 |
| Bio-cell anode bias voltage (V) | $U_{\text{cell 2, anode, bias}}$ | 0.3 [Liu2016a] | 0.2 [Ueki2018a] | 0.3 | 0.2 |
| Bio-cell cathode std. potential (V) | $U_{\text{cell 2, cathode, 0}}$ | 0.82 | | | |
| Bio-cell cathode bias voltage (V) | $U_{\text{cell 2, cathode, bias}}$ | 0.47 | | | |
| Bio-cell voltage (V) | $\Delta U_{\text{cell 2}}$ | 2 [Liu2016a] | 1.59 | 2 | 1.59 |
| Bio-cell Faradaic efficiency | $\zeta_{I2}$ | 1.0 | | | |
| **Cellular Electron Transport Parameters** |  |  |  |  |  |
| Membrane potential difference (mV) | $\Delta U_{\text{membrane}}$ | 140 | | 140 | |
| Terminal $e^-$ acceptor potential (V) | $U_{\text{Acceptor}}$ | 0.82 | | | |
| Quinone potential (V) | $U_Q$ | -0.0885 [Bird2011a] | | -0.0885 [Bird2011a] | |
| Mtr EET complex potential (V) | $U_{\text{Mtr}}$ | N/A | -0.1 [Salimijazi2020b] | N/A | -0.1 [Salimijazi2020b] |
| No. protons pumped per $e^-$ | $p_{\text{out}}$ | Unlimited | | Unlimited | |
| **Product Synthesis Parameters** |  |  |  |  |  |
| No. ATPs for product synthesis | $v_{p, \text{ ATP}}$ | [Sheppard2022b] | | | |
| No. NAD(P)H for product | $v_{p, \text{ NADH}}$ | [Sheppard2022b] | | | |
| No. Fd<sub>red</sub> for product | $v_{p, \text{ Fd}}$ | [Sheppard2022b] | | | |
| Product energy density (J molecule<sup>-1</sup>) | $E_{\text{HC}}$ | See **Table S1** | | | |

**Table 2.** Generalized enzymatic reactions for synthesis of straight chain alkane jet fuel components with length *n*, from metabolic intermediates using type II fatty acid synthesis. Reactions for Type II Fatty Acid Synthesis are taken from the Kyoto Encyclopedia of Genes and Genomes (KEGG)^45-47^. Schematic of these reactions is shown in **Fig. 2**. Explicit reactions for synthesis of the alkanes pentane (C_5_) to heptadecane (C_16_) are shown in the computer code supporting this work^51^. FAP: Fatty Acid Photodecarboxylase; ADO: Aldehyde Deformolating Oxygenase; FAH: Fatty-Acid Hydrolase.

| No. | Reaction | EC | KEGG | Comment |
| --- | --- | --- | --- | --- |
| **Initiation** |  |  |  |  |
| 0 | $\text{CO}_2 + \text{H}_2\text{O} \rightarrow \text{HCO}_3^- + \text{H}^+$ | 4.2.1.1 | R10092 | |
| 1 | $\text{ATP} + \text{acetyl-CoA} + \text{HCO}_3^- \rightarrow \text{ADP} + \text{orthophosphate} + \text{malonyl-CoA}$ | 6.4.1.2 | R00742 | |
| 2 | $\text{malonyl-CoA} + \text{H-[acp]} \rightarrow \text{malonyl-[acp]} + \text{H-CoA}$ | 2.3.1.39 | R01626 | |
| 3 | $\text{acetyl-CoA} + \text{H-[acp]} \rightarrow \text{acetyl-[acp]} + \text{H-CoA}$ | 2.3.1.86 | R01624 | |
| 4A | $\text{acetyl-[acp]} + \text{malonyl-[acp]} \rightarrow \text{acetoacetyl-[acp]} + \text{CO}_2 + \text{H-[acp]}$ | 2.3.1.41 | R04355 | Even chain length fatty acid initiation. |
| 4B | $\text{propionyl-[acp]} + \text{malonyl-[acp]} \rightarrow \text{pentanyl-[acp]} + \text{CO}_2 + \text{H-[acp]}$ | 2.3.1.41 | R04355 | Odd chain length fatty acid initiation. |
| **Extension** |  |  |  |  |
| 5 | $3\text{-oxo-}n\text{-anoyl-[acp]} + \text{NADPH} + \text{H}^+ \rightarrow (\text{R})\text{-3-hydroxy-}n\text{-enoyl-[acp]} + \text{NADP}^+$ | 1.1.1.100 | R02767 | |
| 6 | $(\text{R})\text{-3-hydroxy-}n\text{-enoyl-[acp]} \rightarrow \text{trans-}n\text{-2-enoyl-[acp]} + \text{H}_2\text{O}$ | 4.2.1.59 | R04428 | |
| 7 | $\text{trans-}n\text{-2-enoyl-[acp]} + \text{NADPH} + \text{H}^+ \rightarrow n\text{-anoyl-[acp]} + \text{NADP}^+$ | 1.3.1.9 | R04429 | |
| 8 | $n\text{-anoyl-[acp]} + \text{malonyl-[acp]} \rightarrow 3\text{-oxo-}(n+2)\text{-anoyl-[acp]} + \text{CO}_2 + \text{H-[acp]}$ | 2.3.1.41 | R04355 | Fatty acid extension reaction. |
| **Termination** |  |  |  |  |
| 9A | $n\text{-anoyl-[acp]} + \text{NADPH} + \text{H}^+ \rightarrow \text{H-[acp]} + \text{NADP}^+ + n\text{-aldehyde}$ | 1.2.1.80 | R09484 | ADO pathway option for termination. |
| 10A | $n\text{-aldehyde} + \text{O}_2 + 2 \text{NADPH} + 2 \text{H}^+ \rightarrow (n-1)\text{-ane} + \text{formate} + \text{H}_2\text{O} + 2 \text{NADP}^+$ | 4.1.99.5 | R03728 | ADO pathway option for termination. |
| 9B | $n\text{-anoyl-[acp]} + \text{H}_2\text{O} \rightarrow \text{H-[acp]} + n\text{-anoic acid}$ | 3.1.2.14 | R04014 | FAP pathway option for termination with FAH enzyme |
| 10B | $n\text{-anoic acid} + \text{photon} \rightarrow (n-1)\text{-ane} + \text{CO}_2$ | 4.1.1.106 | R11915 | FAP pathway option for termination with FAP enzyme. |
| 11 | $\text{formate} + \text{NAD}^+ \rightarrow \text{CO}_2 + \text{NADH}$ | 1.17.1.9 | R00519 | |
| 12 | $\text{NADH} + \text{NADP}^+ \rightarrow \text{NADPH} + \text{NAD}^+$ | 1.6.99.1 | R00282 | |

In this work, we consider the electrical energy cost and energy conversion efficiency of electromicrobial production of a model drop-in compatible jet fuel blend consisting of 85% (by number of molecules) straight-chain alkanes and 15% terpenoids that can be used without significant addition of petroleum-based molecules. That being said, it is expected that any biofuel for aviation will need to be supplemented with non-fuel additives which change the physical characteristics of the fuel in order to meet stringent international standards^31^. These additives generally serve as antioxidants, metal deactivators, lubricity improvers, static dissipators, biocides, icing inhibitors, and thermal stabilizers.A full list of chemical formulas, molecular weights and energy densities of products considered in this article is shown in **Table S1**.

As opposed to physiochemical fuel generation processes like Fischer-Tropsch, the output of microbial production can be specifically tailored through genetic engineering. This could be uniquely suited to the production of highly standardized fuels such as those required in aviation. By combining advances in metabolic engineering with EMP, it should be possible to produce drop-in fuels which do not require the agricultural inputs necessitated by earlier generations of biofuels. In this work, we calculate the maximum efficiency of an EMP-to-Jet process, using either EEU or H_2_-oxidation as a source of reducing power.

### Theory

Here, we expand on our prior work^11,21,32^ for the prediction of upper limit efficiency for a highly engineered organism which produces hydrocarbons for the express purpose of creating a drop-in jet fuel. A full set of symbols for this article are shown in **Table S2**.

Within this study, we calculate the efficiency of hydrocarbon production utilizing a set of model parameters shown in **Table 1**. We assume access to a reservoir of atmospheric CO_2_ either for *in vivo* production (**Fig. 1A**) or electrochemical reduction of CO_2_ (**Fig. 1B**), (*e*.*g*., to formate^33^). Bio-electrical energy is realized either by H_2_ reduction and diffusion (**Fig. 1C** part 1) or delivery of electrons to the cell either via an diffusible intermediary (*e*.*g*., water soluble quinones like anthra(hydra)quinone-2,6-disulfonate (AHDS_red_/AQDS_ox_)^14^) or through direct electrical contact with an anode^14,34^ (**Fig. 1C** part 2) (both processes are known as Extracellular Electron Uptake or EEU). In both cases, the electrode provides the microbe with reducing power enabling regeneration of the intracellular reductants NAD(P)H and reduced Ferredoxin (Fd_red_), and the energy carrying molecule ATP (**Fig. 1C**). Microbial maintenance energy is assumed to be negligible for this maximal efficiency calculation, and thus all electrical energy utilized generates a hydrocarbon with internal energy *E*_HC_, at a rate of 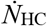, with the microbial cell persisting as a ‘bag of enzymes’^11,35^.

The energy conversion efficiency of electricity to product, *η*_EP_, is calculated from the ratio of the amount of chemical energy stored per second 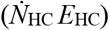, relative the the power input to the system, *P*_e, T_,

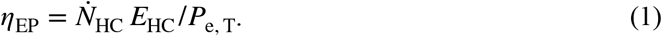

Additionally, the energy required to generate one mole of desired hydrocarbon product, *L*_EP_, is calculated from the following,

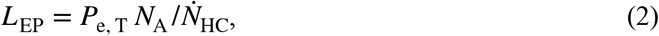

where *N*_A_ is Avogadro’s number.

In the case of *in vivo* carbon fixation (**Fig. 1A**), the upper limit of electrical-to-chemical conversion efficiency is equivalent to the energy density of the hydrocarbon relative to the required charge to synthesize it from CO_2_ and the potential difference across the bio-electrochemical cell^11^,

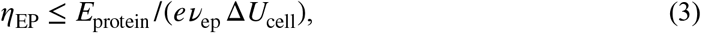

where, *e* is the fundamental charge, *ν*_ep_ is the number of electrons required for synthesis, and Δ*U*_cell_ is the potential difference across the bio-electrochemical cell. Therefore the input electricity required for a mole of hydrocarbon is,

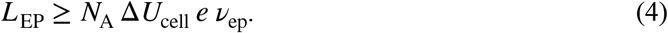

In the case of initial electrochemical reduction of CO_2_ (**Fig. 1B**), we substitute *ν*_ep_ for a more complex function, *ν*_e, add_, which describes the number of electrons required to fully reduce an electrochemical reduction product (almost certainly a C_1_-compound like formate) to the final product^11^,

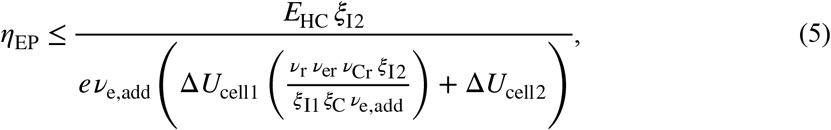

where *ν*_r_ is the required number of primary reduction products needed to generate a final product, *ν*_er_ is the number of electrons to reduce CO_2_ to this primary product (*e*.*g*., 2 in the case of formate), *ν*_Cr_ is the number of carbon atoms per primary fixation product (*e*.*g*., 1 in the case of formate), *ξ*^I2^ is the Faradaic efficiency of the bio-electrochemical cell, *ξ*^I1^ is the Faradaic efficiency of the primary abiotic cell 1, and *ξ*_C_ is the carbon transfer efficiency from cell 1 to cell 2.

Herein the required electrical energy to produce a unit-mole of hydrocarbon via electrochemical CO_2_ reduction is^21^,

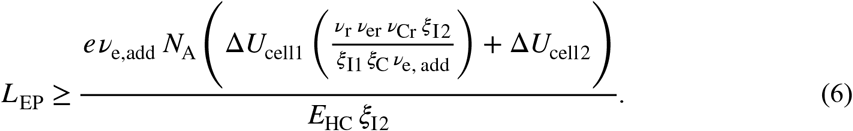

We calculate the number of electrons needed for product synthesis (*ν*_ep_ or *ν*_e, add_) from a model of electron uptake by H_2_-oxidation or EEU, and from the number of NAD(P)H (for the purposes of this analysis we consider NADH and NADPH as equivalent given their identical redox potentials), reduced Ferredoxin, and ATP required by the metabolic pathway used for hydrocarbon synthesis^11^. For electron delivery by

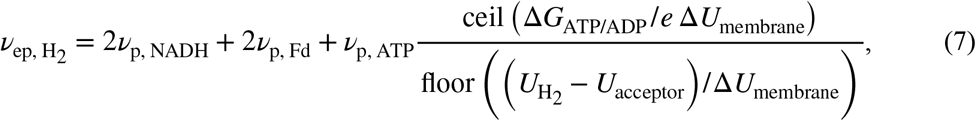

wherein Δ*G*_ATP/ADP_ is the free energy required for the regeneration of ATP, Δ*U*_membrane_ is the potential difference across the cell’s inner membrane due to the proton gradient, *U*_H2_ is the standard redox potential of proton reduction to H_2_, *U*_acceptor_ is the potential of the electron acceptor, *U*_NADH_ is the potential of NADH, and *U*_Fd_ is the potential of Ferredoxin. The ceil function rounds up to the nearest integer, while the floor function rounds down to the nearest integer.

Furthermore, for electron delivery by EEU^11^,

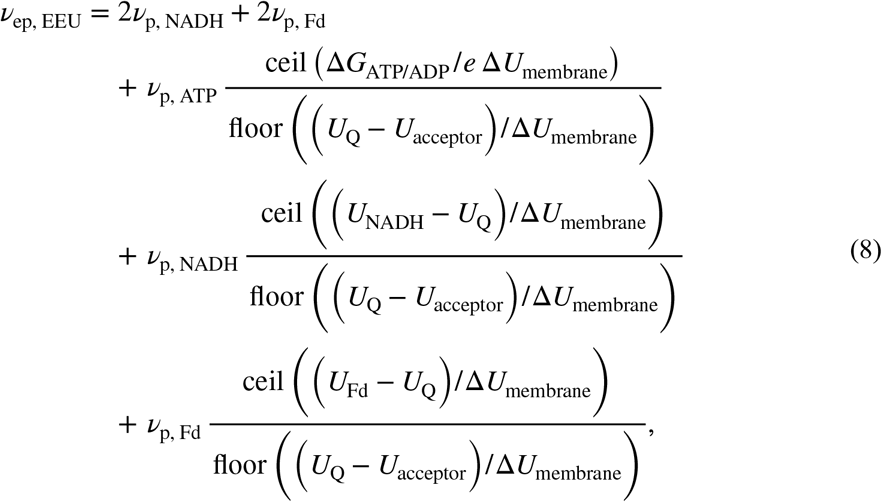

wherein, *U*_Q_ is the redox potential of the quinone inner membrane electron carrier. An expansive version of this derivation can be found in the supplement to our previous work by Salimijazi *et al*.^11^.

In this study, several carbon fixation/assimilation pathways are compared, with the Calvin Cycle (CBB) being the primary focus of our analysis. Other pathways compared include the 3-hydroxypropionate/4-hydroxybutyrate (3HP/4HB) Pathway^36,37^, the 3-hydroxypropionate (3HP) Pathway^38,39^ the Dicarboxylate/ 4-hydroxybutyrate (4HB) Pathway^40^, the Wood-Ljungdahl (WL) Pathway^41^, the Formolose (FORM) Assimilation Pathway^17^, and the Reductive Tricarboxylic Acid (rTCA) pathway^42^. Enzymatic reactions for the production of the metabolic intermediate acetyl-CoA from CO_2_ or formate are shown in **Table S3**. Additionally, we explore two different alkane termination pathways: the ADO (Aldehyde Deformolating Oxygenase) pathway^43^ (**Fig. 2** and reactions 9A and 10A in **Table 2**), and the FAP (Fatty Acid Photodecarboxylase) pathway^44^ (**Fig. 2** and reactions 9B and 10B in **Table 2**).

NAD(P)H, reduced Ferredoxin, and ATP requirements for synthesis of straight-chain alkanes and terpenoids from acetyl-CoA were found by flux balance analysis of the set of biochemical reactions used in their synthesis. Pathways for production of individual straight-chain alkane compounds are compiled in **Fig. 2** and **Table 2** from listings of reactions of Type II Fatty Acid Synthesis in the Kyoto Encyclopedia of Genes and Genomes (KEGG)^45−47^. Pathways for the production of the terpenoid compounds were compiled in an earlier review article^48^ using information from Cheon *et al*.^49^ with additional pathways from Peralta-Yahya *et al*.^50^ and are shown in **Fig. 3** and **Table 3**.

**Table 3.** Reactions for synthesis of aromatic components of jet fuel. Reaction numbers (column 1) correspond to reactions shown in **Figure 2**. Pathways for the production of the terpenoid compounds were compiled by Adesina *et al*.^48^ using information from Cheon *et al*.^49^ with additional pathways from Peralta-Yahya *et al*.^50^

| # | Reaction | EC | KEGG |
| --- | --- | --- | --- |
| 1 | $2 \text{ acetyl-CoA} \rightarrow \text{CoA} + \text{acetoacetyl-CoA}$ | 2.3.1.9 | R00238 |
| 2 | $\text{acetyl-CoA} + \text{H}_2\text{O} + \text{acetoacetyl-CoA} \rightarrow (\text{S})\text{-3-Hydroxy-3-methylglutaryl-CoA} + \text{CoA}$ | 2.3.3.10 | R01978 |
| 3 | $(\text{S})\text{-3-Hydroxy-3-methylglutaryl-CoA} + 2 \text{ NADPH} + 2 \text{ H}^+ \rightarrow (\text{R})\text{-Mevalonate} + \text{CoA} + 2 \text{ NADP}^+$ | 1.1.1.34 | R02082 |
| 4 | $\text{ATP} + (\text{R})\text{-mevalonate} \rightarrow \text{ADP} + (\text{R})\text{-5-Phosphomevalonate}$ | 2.7.1.36 | R02245 |
| 5 | $\text{ATP} + (\text{R})\text{-5-phosphomevalonate} \rightarrow \text{ADP} + (\text{R})\text{-5-diphosphomevalonate}$ | 2.7.4.2 | R03245 |
| 6 | $\text{ATP} + (\text{R})\text{-5-diphosphomevalonate} \rightarrow \text{ADP} + \text{Orthophosphate} + \text{isopentenyl diphosphate} + \text{CO}_2$ | 4.1.1.33 | R01121 |
| 7 | $\text{isopentenyl diphosphate} \rightarrow \text{dimethylallyl diphosphate}$ | 5.3.3.2 | R01123 |
| 8 | $\text{dimethylallyl diphosphate} + \text{isopentenyl diphosphate} \rightarrow \text{diphosphate} + \text{geranyl diphosphate}$ | 2.5.1.1 | R01658 |
| 9 | $\text{Geranyl diphosphate} \rightarrow (-)\text{-alpha-pinene} + \text{diphosphate}$ | 4.2.3.119 | R05765 |
| 10 | $\text{geranyl-diphosphate} + \text{H}_2\text{O} \rightarrow \text{geraniol} + \text{diphosphate}$ | 3.1.7.11 | R08396 |
| 11 | $\text{geranyl-diphosphate} \rightarrow (\text{S})\text{-limonene} + \text{diphosphate}$ | 4.2.3.16 | R02013 |
| 12 | $\text{geranyl-diphosphate} + \text{isopentenyl diphosphate} \rightarrow \text{diphosphate} + (\text{2E,6E})\text{-farnesyl-diphosphate}$ | 2.5.1.10 | R02003 |
| 13 | $(\text{2E,6E})\text{-farnesyl-diphosphate} \rightarrow \text{alpha-farnesene} + \text{diphosphate}$ | 4.2.3.46 | R08696 |
| 14 | $(\text{2E,6E})\text{-farnesyl-diphosphate} \rightarrow (\text{E})\text{-alpha-bisabolene} + \text{diphosphate}$ | 4.2.3.38 | R08370 |

The set of reactions for product synthesis are balanced in a custom flux balance analysis code that calculates the required NAD(P)H, Fd_red_, ATP, CO_2_, and formate inputs needed to produce a single target molecule^21^. In this, a stoichiometric matrix is generated wherein the vector 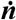 encodes the change in number of molecules over a single round of the reaction cycle; **S**_**p**_ is a matrix that encodes the stoichiometry for each reaction in the pathway; and ***ν*** is a flux vector describing the number of times each reaction is utilized throughout a reaction cycle,

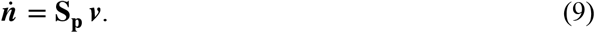

In this framework, 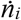 is constrained to be zero for each reactant designated as an intermediate (*e*.*g*., acetyl-CoA, acetoacetyl-CoA, *etc*.; but not CO_2_, NAD(P)H, Fdred, or ATP) in the reaction network (*i*.*e*., its number should not grow or shrink over an entire reaction cycle, leaving the host cell’s chemical state unchanged). The optimized stoichiometry for synthesis of each hydrocarbon is enumerated in the supporting online code repository for this article^51^.

A second program^11^ then takes the required NAD(P)H, Fd_red_, and ATP to produce all target molecules and calculates the required input energy to generate these compounds based on their energy density and molecular weight. We use the results of this calculation to determine the most efficient method of electron delivery (H_2_-oxidation or EEU); methods of carbon-fixation or carbon-assimilation; and fatty-acid synthesis cycle termination reaction are most efficient and economically viable in practice for synthesis.

## Results and Discussion

### Electromicrobial Production Synthesizes Individual Straight-chain Alkanes with ≈ 40% Efficiency

We calculated the energy requirements for synthesis of one mole of the straight-chain alkane (aka paraffinic) compounds pentane (C_5_) to heptadecane (C_16_) using the Calvin CO_2_-fixation cycle (CBB) with H_2_-oxidation and EEU for electron uptake (**Fig. 4A**). The corresponding electrical-to-chemical energy conversion efficiencies for production of these compounds are shown in **Fig. 4B**.

**Figure 4.**
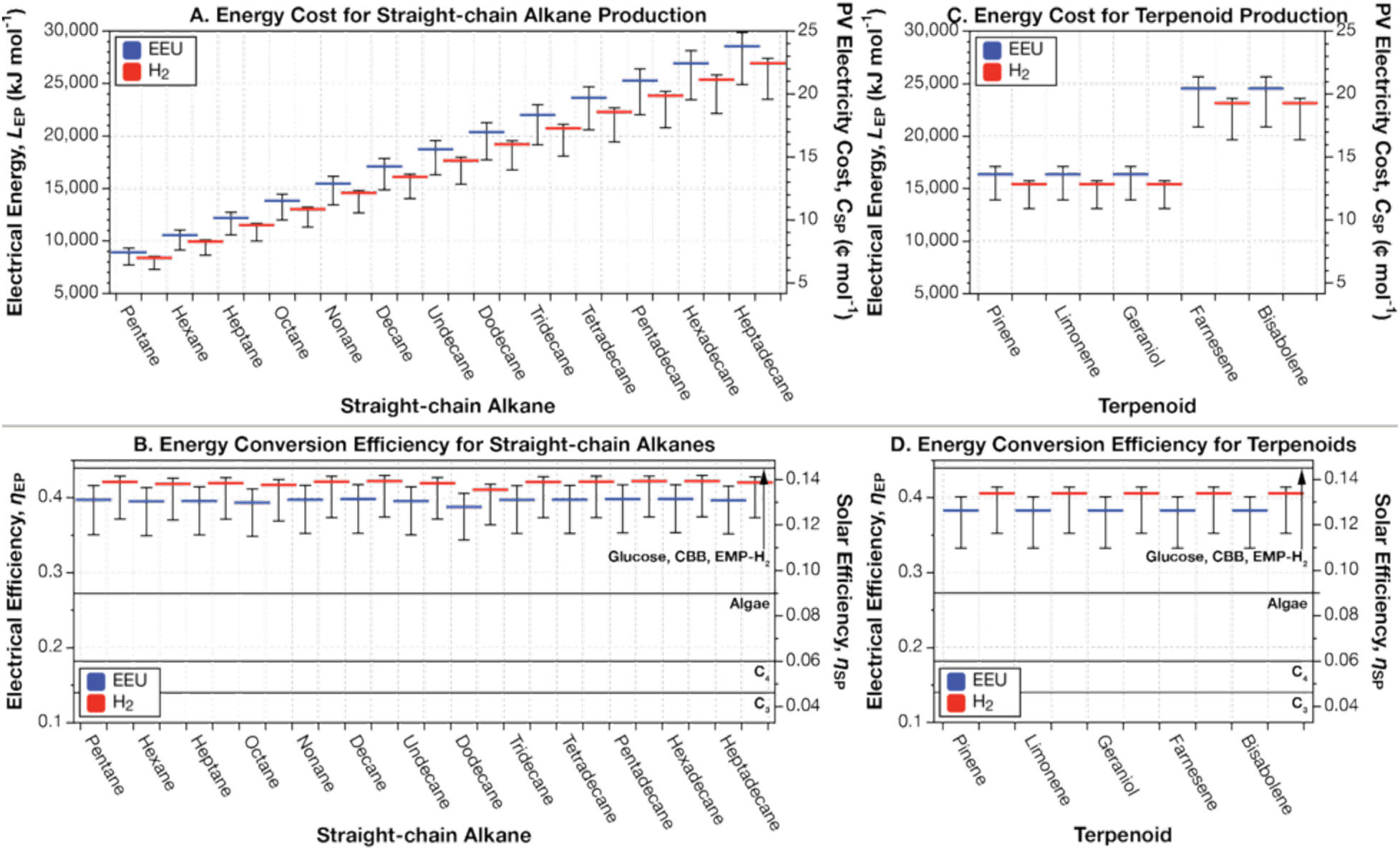
Electrical energy requirements and energy conversion efficiencies for straight-chain alkane and terpenoid production. (**A**) Energy input for, and (**B**) energy conversion efficiency of straight-chain fatty alkane biosynthesis using the Calvin CO_2_-fixation cycle with the ADO alkane termination pathway. (**C**) Energy input required for, and (**D**) energy conversion efficiency of terpenoid compound biosynthesis. A sensitivity analysis by Salimijazi *et al*.^11^ found that the biggest source of uncertainty in the energy input and efficiency calculation is the potential difference across the inner membrane of the cell (Δ*U*_membrane_). Estimates for the trans-membrane voltage range from 80 mV (BioNumber ID (BNID) 10408284 (ref. 70) to 270 mV (BNID 107135), with a most likely value of 140 mV (BNIDs 109774, 103386, and 109775). The central value (thick blue or red bar) corresponds to the most likely value of the trans-membrane voltage of 140 mV. Our sensitivity analysis found that Δ*U*_membrane_ = 280 mV produces lower efficiencies (hence a higher energy input), while Δ*U*_membrane_ = 80 mV produces higher efficiencies (and hence lower energy inputs)^11^. The right axis in panels **A** and **C** shows the minimum cost of that solar electricity, assuming that the United States Department of Energy’s cost target of 3 ¢ per kWh by 2030 can be achieved^71^. The right axes in panels **B** and **D** show the solar-to-product energy conversion efficiency, assuming the system is supplied by a perfectly efficient single-junction Si solar photovoltaic (solar to electrical efficiency of 32.9%^72^. For comparison, we have marked the upper limit solar-to-biomass energy conversion efficiencies of C_3_, C_4_ (refs. 73,74), algal photosynthesis^75^, and upper limit electromicrobial production conversion efficiency of glucose^21^ using H_2_-oxidation and the Calvin cycle on the right axes of panels **B** and **D**. This figure can be reproduced by running the codes Info-Fig4a&c.py, Info-Fig4b&d.py, Generate-Fig4a&c.py, and Generate-Fig4b&d.py in the EMP-to-Jet online code repository^51^.

Each cycle of the Type II Fatty Acid Synthesis pathway adds an additional two carbon atoms to the fatty acid chain, requiring additional CO_2_ fixation as well as an additional 3 NADH to attach the carbon to the growing fatty acid and to add two protons to the chain.

For electron delivery with H_2_, the energy cost of production of paraffinic compounds rises approximately linearly from 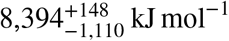 for pentane to 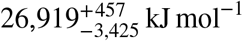 for heptadecane (**Fig. 4A** red bars). Each additional carbon added demands an additional 1,544 kJ mol^−1^ of electrical energy.

While energy costs for the synthesis of straight-chain alkane compounds rise linearly, so does their heat of combustion (**Table S1**). This means that the electrical energy conversion efficiency using H_2_-oxidation and the Calvin cycle for all alkane compounds remains constant at ≈ 41.6% (**Fig. 4B** red bars). There is small but noticeable drop in efficiency for dodecane due to the lower than expected reported heat of combustion (**Table S1**). The energy conversion efficiency of straight-chain alkanes is comparable to that for butanol (44.0% when using the Calvin cycle and H_2_-oxidation^11,21^) and glucose (44.6% when using the Calvin cycle and H_2_-oxidation^11,21^).

### Electromicrobial Production Synthesizes Individual Terpenoid Compounds with ≈ 40% Efficiency

The energy cost of production terpenic compounds is 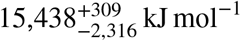 for pinene, limonene and geraniol and 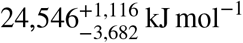 for farnesene and bisabolene when using electron delivery by H_2_-oxidation and the Calvin cycle for CO_2_-fixation (**Fig. 4C** red bars). The energy consumption for production of pinene, limonene, and geraniol are identical, due to the similar chemical composition of these molecules and lack of energy consuming reactions needed for the differentiating steps in their production. While farnesene and bisabolene are also produced by the terpenoid pathway, these compounds require additional energy-consuming steps in their synthesis (**Fig. 3**).

As is the case with the straight-chain alkane compounds, the electrical to chemical energy conversion efficiency for terpenic compounds are identical (40.5% for H_2_-oxidation and the Calvin cycle), resulting from the very similar combustion energies (**Fig. 4D** red bars).

### Electron Uptake by H_2_-Oxidation Produces the Higher Energy Conversion Efficiencies than EEU

It is clear from all panels in **Fig. 4** that electron delivery by H_2_-oxidation (**Fig. 4** red bars) results in lower energy costs and higher efficiencies for synthesis of a mole of any jet-fuel component than EEU (**Fig. 4** blue bars). As we have discussed in earlier articles^11,21,32^ the small difference in efficiency is due to the fact that electrons in EEU-mediated EMP are delivered to the cell at a higher redox potential than NADH, requiring the use of reverse electron transport to raise their energy sufficiently to reduce NADH^11,14,34^ (−0.32 V vs. Standard Hydrogen Electrode (SHE) for NADH, while the redox potential of the electron-accepting Mtr complex is ≈ -0.1 V vs. SHE^52^). In contrast, H_2_-mediated EMP delivers electrons at a redox potential slightly lower than NADH (−0.42 Volts vs. Standard Hydrogen Electrode for H_2_ vs. -0.32 V for NAD(P)H) meaning that that direct reduction of NADH is possible.

The difference in energy input between H_2_- and EEU-mediated EMP grows from 504 kJ mol^−1^ for pentane to 1,615 kJ mol−1 for heptadecane (**Fig 3A**). Despite this, the difference in energy conversion efficiency between H_2_- and EEU-mediated production of straight-chain alkanes remains constant at 2.3% (**Fig. 4B**).

The difference in energy input between H_2_- and EEU-mediated EMP of the terpenoids pinene, limonene an geraniol is 926 kJ mol^−1^ (**Fig. 4C**). The energy difference between H_2_ and EEU-mediated EMP for farnesene and bisabolene rises to 1,389 kJ mol^-1^. The efficiency difference between synthesis of terpenoids by H_2_- and EEU-mediated EMP remains constant at 2.3% (the same as for straight-chain alkanes) (**Fig. 4D**).

### Efficiency of EMP-to-Jet Can be Raised to 50% by Adopting More Efficient CO_2_-fixation Pathways

To further explore possible efficiency improvements in our system, we compared the effect of changing carbon fixation methods on the efficiency of EMP of a blend of alkanes and terpenoids (85% total molarity straight-chain alkanes from C_10_ to C_16_, 15% of each alkane; 15% molarity terpenoids, 3% of each terpenoid) (**Figs. 5A** and **B**). We calculated the number of NAD(P)H, Fd_red_, and ATP needed for synthesis of each component of the blend and then calculated their weighted averaged (depending on the amount of molecules of each component in the blend). The weighted average requirements for NAD(P)H, Fd_red_, and ATP were input into our code for calculating the energy requirements and efficiency of EMP (a similar approach was used by Wise *et al*.^21^ to calculate the energy requirements and efficiency of EMP of a blend of amino acids).

**Figure 5.**
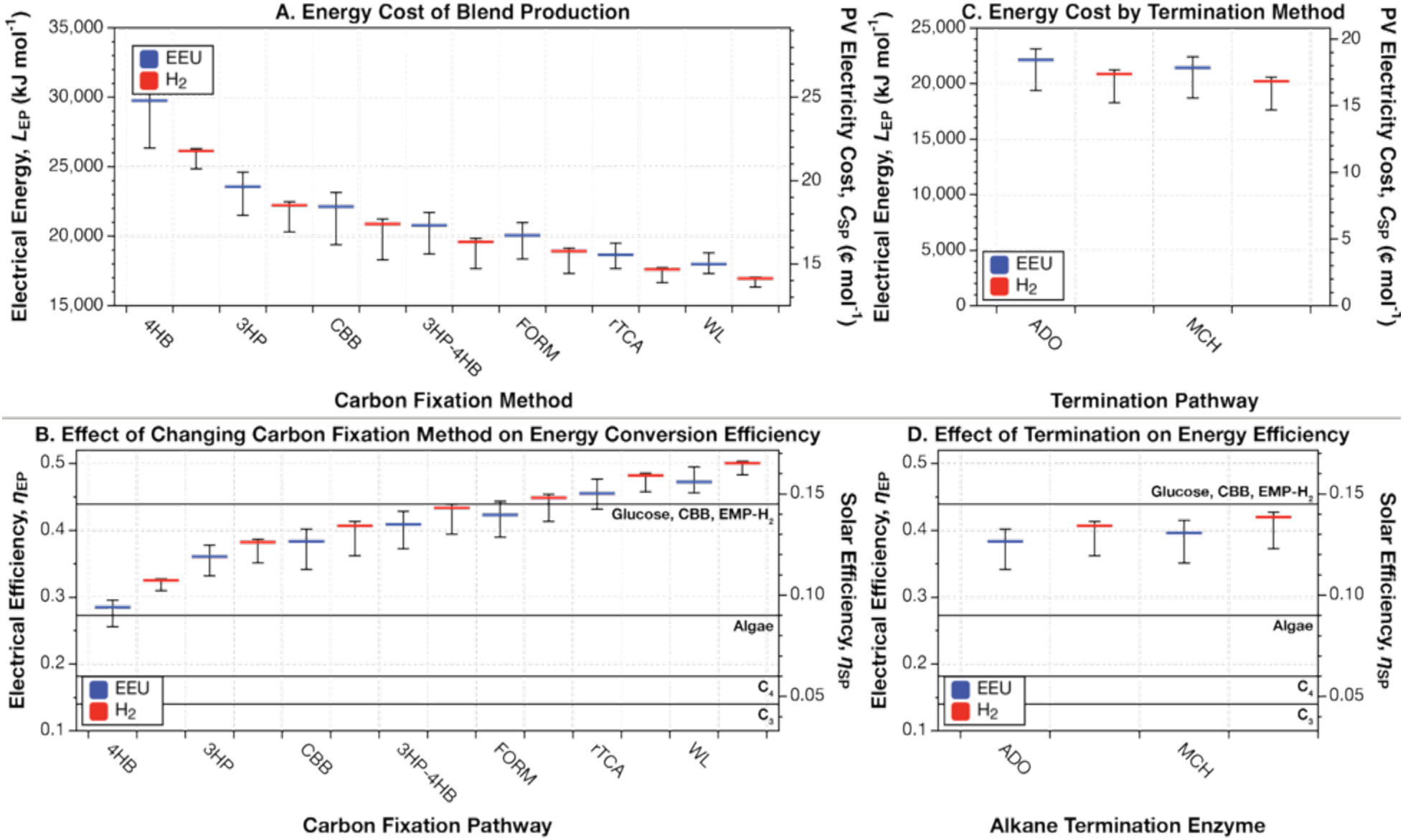
Effect of changing carbon-fixation (or assimilation) method and alkane termination pathway on the energy requirements and conversion efficiency of a model jet fuel blend. Effect of changing carbon-fixation method on (**A**) energy input for, and (**B**) energy conversion efficiency of production of a model jet fuel blend containing 85% molarity (equal numbers of C_10_ to C_16_ molecules) and 15% terpenoids (equal numbers of each of the 5 terpenoids) when using the ADO alkane termination pathway. Effect of changing alkane termination method on (**C**) energy input for, and (**D**) energy conversion efficiency of production of model jet fuel blend when using the Calvin CO_2_-fixation cycle. The central value (thick blue or red bar) corresponds to the most likely value of the trans-membrane (Δ*U*_membrane_) voltage of 140 mV. Meanwhile, Δ*U*_membrane_ = 280 mV produces lower efficiencies (hence a higher energy input), while Δ*U*_membrane_ = 80 mV produces higher efficiencies (and hence lower energy inputs)^11^. This figure can be reproduced by running the codes Info-Fig5a&c.py, Info-Fig5b&d.py, Generate-Fig5a&c.py, and Generate-Fig5b&d.py in the EMP-to-Jet online code repository^51^.

The right choice of carbon fixation mechanism can reduce energy costs for production of a mole of the jet fuel blend by more than a third. For electron delivery by H_2_-oxidation, the energy cost of production of the blend decreases from 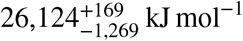 when using the 4HB pathway to 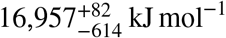 when using the WL pathway (a 35% reduction). Likewise, when using EEU for electron delivery, the energy cost drops from 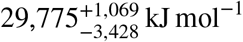 to 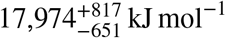 (a 40% reduction).

The efficiency of jet fuel blend production can be raised to 50%. When using EEU for electron delivery, the efficiency of jet fuel blend production rises from 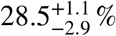 when using the 4HB pathway, to 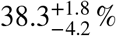 when using the Calvin cycle (CBB), and to 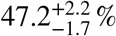 when using the WL pathway. When using H_2_-oxidation, the efficiency of the jet fuel blend production rises from 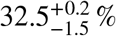 when using the 4HB pathway, to 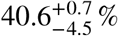 when using CBB, and to 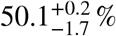 when using the WL pathway.

### Choice of Alkane Termination Reaction Can Increases Efficiency of Jet Fuel Blend Production by ≈ 2%

An ADO pathway for termination has been widely explored in relation to hydrocarbon chain termination in *E. coli*^53^ (**Fig. 2**, and **Table 2** reactions 9A and 10A). This pathway catalyzes an initial acyl-[acp] replacement reaction, substituting the complex for a hydrogen, expending an NADH in the process and forming an aldehyde. This aldehyde is then cleaved, removing the carbonyl group and substituting a hydrogen in its place, forming a fatty acid and expending two NADH in the process, cleaving an O_2_ molecule in the process. This pathway has the benefit of being native to many cyanobacteria and has been shown to be functional in hydrocarbon production and is very well studied and explored. However, it is a slow process (1 min^-1^) without modification^54-56^, and expends considerable excess energy to cleave a single carbon.

The FAP pathway is less familiar, with its greatest activity associated with the membrane of chloroplasts in algal cells^57^. This pathway initially hydrolyzes the acyl-[acp] group, replacing it with a terminal carboxyl group. Then the photoactive decarboxylase FAP enzyme cleaves the carboxyl group with the input of blue light. This pathway has the benefit of being more energy efficient, requiring less chemical energy to achieve and requires an input of further electrical energy which can be generated with ease. However, the activity of this pathway in heterologous hosts has barely been studied beyond a handful of studies in *E. coli*^58,59^.

These results indicate what would be expected of these paths, wherein the FAAP/ADO pathway utilizes three more NADH per straight-chain alkane molecule produced, which decreases the overall efficiency as the resulting compounds are not any more energy dense. Production of the jet fuel blend with the ADO pathway and the Calvin cycle costs 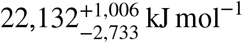 when using EEU and 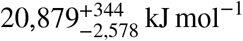 when using H_2_. By contrast, swapping the termination method to the MCH pathway reduces energy costs to 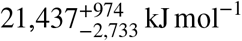 when using EEU and 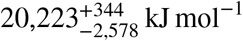 when using H_2_ (**Fig. 5C**). These cuts in energy costs increase the efficiency of blend production by ≈ 2% (**Fig. 5D**). Notably, we assume here that blue light is free, and the energy associated with the utilization of a constant blue light source has a cost not represented in this figure, and thus may alter the results.

### In Almost All Cases, Efficiency of EMP-to-Jet is Higher than Conventional Biofuels

How do the upper-limit efficiencies predicted for EMP-to-Jet production compare with the production efficiencies of existing biofuel technologies? Many studies of the efficiency of biofuel production concern biomass-to-fuel, and biomass-to-wheel energy conversion efficiency^60^. Traditional energy input estimates of biofuels are not wrong. Quite rightly, sunlight has been thought of as free of cost and global warming concerns. Furthermore, traditional analyses rightly concern themselves with necessary fossil energy inputs. However, if global agricultural production expands to simultaneously enable large scale biofuel production and produce larger volumes of food, this could cause significant competition between land for crops and land for wilderness^61^. As a result, land for agriculture could become an increasingly scarce commodity, making efficient of use of sunlight increasingly important^19,21,62^.

What effect could widespread biofuel production have on land use? To estimate the minimum amount of land that we would need to be converted to biofuel crop production, *A*_land_, Brenner *et al*.^62^ compared average transportation power demand, *P*_transport_, with the amount of solar power that is captured as biomass and converted to biofuel per unit area, *P*_fuel_,

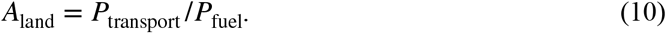

The average transportation power can be estimated by summing all transportation energy uses in a given period of time, *τ* (*i*.*e*., a year),

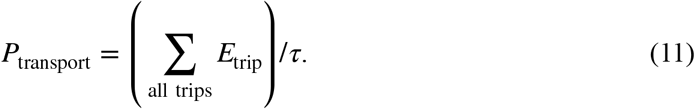

For instance, averaged over a year, the power use of all transportation in the United States is ≈ 1 terawatt (TW)^62^. The US Energy Information Administration estimates global demand for just aviation fuels will reach ≈ 30 quadrillion British Thermal Units per year by 2050 (1 TW).

The amount of solar power captured as biomass and converted to fuel can be estimated from the average available solar power, ⟨*P*_⊙_⟩; the efficiency of photosynthetic conversion of solar to biomass energy, *η*_SB_; and the conversion of efficiency of biomass to fuel, *η*_BF_. Thus,

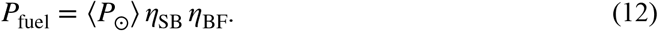

What land area needed to produce transportation fuel at the rate of 1 TW? Assuming a day/night averaged solar power at the mid-latitudes (between the tropics and the polar circles) of ≈ 200 W m^-2^ (ref 62), the total amount of land needed to capture solar power at the rate *P*_transport_ is,

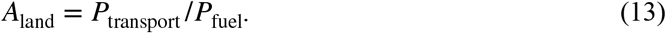

Assuming 100% conversion efficiency of biomass to fuel (high, but not so wide of the mark^60^), the total land surface area needed to deliver transportation fuel at a rate of 1 TW, relative to US cropland area of 1.59 × 10^12^ m^2^ (ref 63),

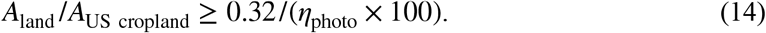

Thus, if photosynthetic efficiency is 1%, at least 32% of US cropland area will be needed to supply transportation fuel at a rate of 1 TW (note that Brenner *et al*. estimated this as 28%, but since their report US cropland area has shrunk by ≈ 50 million acres or 2 × 10^11^ m2 (ref 63)).

On the other hand, if the solar-to-fuel efficiency can be raised to 10% (just below the predicted maximum efficiency of EMP-to-Jet schemes), the required land area can be reduced to only 3.2% of US cropland area, or only 0.7% of combined US crop, forest and grass and pasture land area of 6.80 × 10^12^ m^2^ (ref 63).

On a global scale, the minimum land area needed to generate fuel at a rate of 1 TW, assuming global cropland area of 1.87 × 10^13^ m^2^ (ref. 64),

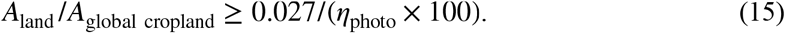

In other words, if photosynthetic efficiency is 1%, then 2.7% of global cropland will need to dedicated to aviation fuel crops. Is 2.7% of global cropland a big deal? The effect of the war in Ukraine on global food supply can give us some sense of the answer. The Russian invasion took ≈ half of Ukraine’s ≈ 50 million tonnes of grain exports off the global market, placing noticeable pressure on global food prices. How much of the world’s food does Ukraine produce?: less than 1% of the world’s annual agricultural output of 9.2 billion tonnes per year. Given this, 2.7% of the world’s cropland is something should not be abused lightly.

## Conclusions

In this article we consider the thermodynamic constraints on the electromicrobial production of a drop-in compatible jet fuel substitute containing 85% C_10_ to C_16_ straight-chain alkanes and 15% terpenoids. We find that the biggest lever for changing production efficiency is the choice of carbon fixation (or assimilation) method used by the microbial production chassis (**Figs. 5A** and **B**). When using H_2_-oxidation for electron delivery, and the widely-studied and used (probably the most widely used enzymatic pathway on Earth) Calvin cycle for carbon-fixation, the production efficiency of the jet fuel blend is 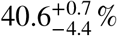. Swapping the Calvin cycle for the 4HB pathway drops the production efficiency to 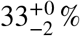, while swapping it for the highly efficient Wood-Ljungdahl pathway raises efficiency to 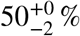. Swapping the alkane termination pathway from the well-known ADO pathway to the less-studied MCH pathway only increases efficiency from 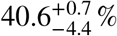 to 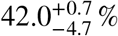 (when using H_2_-oxidation and the Calvin cycle).

How can we leverage the theoretical potential for high efficiency production of jet fuels from electricity and CO_2_ that we calculate in this article? In our earlier studies on the electromicrobial production of butanol^11^, and of amino acids and proteins^21^ we identified the challenge of simultaneously operating the high-efficiency, but O_2_-sensitive rTCA and Wood-Ljungdahl pathways with both the H_2_-oxidation and EEU electron uptake mechanisms that both demand small amounts of O_2_ to generate reducing power^14,34,65^. We identified the need for advances in synthetic sub-cellular compartmentalization that would allow O_2_-sensitive processes to run in the presence of O_2_ (ref. 11). Since the time of writing those earlier articles, Kirst *et al*. have demonstrated the use of a bacterial micro compartment to encapsulate an O_2_-sensitive pyruvate formate lyase, a key enzyme in formate utilization^66^, bringing this vision much closer to reality.

We believe the time is right to start scaling up production of jet fuels with EMP. As we have noted in earlier articles, the efficiency of electromicrobial production using H_2_-oxidation is typically ≈ 2% higher than electromicrobial production using EEU^11,21^. However, scale up H_2_-mediated EMP is complicated by the low solubility of H_2_ in water^11,67^. Simply put, mixing H_2_ into water demands a significant fraction of electrical power that could be put into making a biofuel^11^. However, as the scale of the system rises, relative energy losses drop. For a system consuming ≈ 2 megawatts (MW) of electrical power, energy losses due to mixing drop to less than 5%^11^. Given that a jet fuel production system will probably need to operate at the gigawatt (GW) scale (a 747 in level flight consumes about 140 MW of power^68^), concerns about mixing H_2_ into water become negligible (at least theoretically). This result led us to conclude that efforts to scale up electromicrobial production of jet fuel should start. The results of this article, that the high production efficiencies we earlier saw for butanol translate to jet fuel, further strengthens our belief in this conclusion.

Is there any role left for EMP mediated by extracellular electron uptake (EEU)? We believe there is. First, it’s important to be up front that our lab works on making EEU-mediated EMP a reality, so we have a vested interest in continued support for the development of this technology. In this, light we leave it to the reader to judge our conclusions. Second, we need to acknowledge just how far ahead H_2_-mediated EMP is in terms of technology readiness level compared with EEU-mediated EMP. Furthermore, the theoretical maximum energy conversion efficiency of H_2_-mediated EMP, everything else being equal, is always ≈ 2% higher than EEU-mediated EMP^11^. H_2_-mediated EMP can already achieve a very high fraction of its theoretical maximum efficiency under lab-scale conditions, and this technology has already been spun out of the lab. Meanwhile, we are aware of only one demonstration of biofuel production with EEU-mediated EMP^69^. However, despite our earlier assertion that efficiency losses due to H_2_-mixing become less and less important with increasing power scale, it is not clear how the high lab scale efficiency of H_2_-mediated EMP will translate into achievable commercial scale efficiency. If H_2_-mediated EMP fails to deliver on the promise of its theoretical and lab-scale efficiency, then EEU-mediated EMP has a chance. Finally, while we normally think of H_2_-mediated and EEU-mediated EMP systems belonging inside an electro-bioreactor (like the one in **Fig. 1**) EEU-mediated EMP offers the possibility of creating quantum dot microbe hybrid systems where quantum dots convert light into electron hole pairs that can be transferred directly to a microbe where they can be used to power CO_2_-fixation. This sort of artificial photosynthesis system has the potential to be extremely simple to operate and might offer very high efficiencies.

## End Notes

### Code Availability

All code used in calculations in this article is available at https://github.com/barstowlab/emp-to-jet and is archived on Zenodo^51^.

### Materials & Correspondence

Correspondence and material requests should be addressed to B.B.

### Author Contributions

Conceptualization, T.J.S., D.A.S. and B.B.; Methodology, T.J.S., D.A.S. and B.B.; Investigation, T.J.S. and D.A.S.; Writing - Original Draft, T.J.S. and D.A.S.; Writing - Review and Editing, T.J.S., D.A.S. and B.B.; Resources, B.B.; Supervision, B.B..

## Acknowledgements

This work was supported by a Cornell Energy Systems Institute Postdoctoral Fellowship to D.A.S.; Cornell University startup funds, a Career Award at the Scientific Interface from the Burroughs Welcome Fund, U.S. Department of Energy Biological and Environmental Research grant DE-SC0020179, and a gift from Mary Fernando Conrad and Tony Conrad to B.B..

