## Supplementary Information for "Upper Limit Efficiency Estimates for Electromicrobial Production of Drop-In Jet Fuels"

### Supplementary Information Tables

**Table S1.** Molecular weights and energy densities for straight chain alkanes.

**Table S2.** Symbols used in this article.

**Table S3.** Carbon-fixation and -assimilation, and nitrogen-fixation reactions considered in this article.

**Table S4.** Acetyl-CoA to propionyl-CoA reactions.

| **Hydrocarbon** | **Molecular Weight (Da)** | **Energy Density (kJ mol$^{-1}$)** | **Energy Density (kJ molecule$^{-1}$)** | **Molecular Formula** |
|---|---|---|---|---|
| Pentane | 72.15 | 3,245.00 | $5.87 \times 10^{-18}$ | $C_5H_{12}$ |
| Hexane | 86.175 | 4,163.20 | $6.91 \times 10^{-18}$ | $C_6H_{14}$ |
| Heptane | 100.2 | 4,817.00 | $8 \times 10^{-18}$ | $C_7H_{16}$ |
| Octane | 114.225 | 5,471.62 | $9.02 \times 10^{-18}$ | $C_8H_{18}$ |
| Nonane | 128.25 | 6,125.00 | $1.02 \times 10^{-17}$ | $C_9H_{20}$ |
| Decane | 142.275 | 6,778.29 | $1.13 \times 10^{-17}$ | $C_{10}H_{22}$ |
| Undecane | 156.3 | 7,431.30 | $1.23 \times 10^{-17}$ | $C_{11}H_{24}$ |
| Dodecane | 170.325 | 7,901.74 | $1.31 \times 10^{-17}$ | $C_{12}H_{26}$ |
| Tridecane | 184.35 | 8,741.10 | $1.45 \times 10^{-17}$ | $C_{13}H_{28}$ |
| Tetradecane | 198.375 | 9,464.66 | $1.56 \times 10^{-17}$ | $C_{14}H_{30}$ |
| Pentadecane | 212.4 | 10,049.10 | $1.67 \times 10^{-17}$ | $C_{15}H_{32}$ |
| Hexadecane | 226.425 | 10,699.10 | $1.78 \times 10^{-17}$ | $C_{16}H_{34}$ |
| Pinene | 136.24 | 6,204.87 | $1.04 \times 10^{-17}$ | $C_{10}H_{16}$ |
| Limonene | 136.24 | 6,167.01 | $1.04 \times 10^{-17}$ | $C_{10}H_{16}$ |
| Geraniol | 154.25 | 6,237.72 | $1.04 \times 10^{-17}$ | $C_{10}H_{18}O$ |
| Farnesene | 204.36 | 9,372.45 | $1.47 \times 10^{-17}$ | $C_{15}H_{24}$ |
| Bisabolene | 204.36 | 9,289.02 | $1.47 \times 10^{-17}$ | $C_{15}H_{24}$ |
| Glucose | 180.16 | 2,801.49 | $4.65 \times 10^{-18}$ | $C_6H_{12}O_6$ |
| Butanol | 74.12 | 2,712.79 | $4.50 \times 10^{-18}$ | $C_4H_{10}O$ |

**Table S1.** Molecular weights and energy densities for products considered in this article. Data from NIST database [NIST2022a].

| Symbol | Unit | Description |
|---|---|---|
| $E_{HC}$ | J molecule$^{-1}$ | Energy carried per hydrocarbon molecule. |
| $\dot{N}_{HC}$ | molecule s$^{-1}$ | Hydrocarbon molecules produced per second. |
| $N_A$ | molecule mol$^{-1}$ | Avogadro constant. |
| F | A s mol$^{-1}$ | Faraday constant. |
| $P_{e,\,T}$ | J s$^{-1}$ | Total electrical power input into electromicrobial production system. |
| $L_{EP}$ | kJ mol$^{-1}$ | Electrical energy cost to generate one mole of product. |
| $C_{SP}$ | ¢ mol$^{-1}$ | Minimum solar electricity cost for synthesis of one mole of product at 2030 solar electricity prices. |
| $M_{HC}$ | g mol$^{-1}$ | Molecular weight of hydrocarbon molecule. |
| $\eta_{EP}$ | % | Electrical to product (*e.g.,* hydrocarbon) energy conversion efficiency. |
| $\eta_{SP}$ | % | Solar to product (*e.g.,* hydrocarbon) energy conversion efficiency. |
| $e$ | A s | Fundamental charge. |
| $\nu_{ep}$ | # | Number of electrons needed for synthesis of a product (*e.g.*, hydrocarbon) molecule. |
| $\Delta U_{cell}$ | V | Potential difference across bio-electrochemical cell. |
| $\nu_{e,\,add}$ | # | Number of electrons needed to convert a $C_1$ compound to a hydrocarbon product. |
| $\nu_r$ | # | Number of primary reduction products to make a molecule of final product. |
| $\nu_{er}$ | # | Number of electrons to reduce $CO_2$ to a primary reduction product. |
| $\nu_{Cr}$ | # | Number of carbon atoms per primary reduction product. |
| $\xi_{I2}$ | # | Faradaic efficiency of the bio-electrochemical cell. |
| $\xi_{I1}$ | # | Faradaic efficiency of the primary abiotic cell. |
| $\xi_C$ | # | Carbon transfer efficiency from cell 1 to cell 2. |
| $\nu_{p,\,NADH}$ | # | Number of NAD(P)H molecules needed to make a final product molecule. |
| $\nu_{p,\,Fd}$ | # | Number of Fd molecules needed to make a final product molecule. |
| $\nu_{p,\,ATP}$ | # | Number of ATP molecules needed to make a final product molecule. |
| $\Delta G_{ATP/ADP}$ | J | Free energy for regeneration of ATP. |
| $\Delta U_{membrane}$ | V | Inner membrane potential difference. |
| $U_{H2}$ | V | Standard potential of proton reduction to $H_2$. |
| $U_{acceptor}$ | V | Standard potential of terminal electron acceptor reduction. |
| $U_Q$ | V | Redox potential of the inner membrane electron carrier. |
| $U_{NADH}$ | V | Standard potential of NADH. |
| $U_{Fd}$ | V | Standard potential of Ferredoxin. |

**Table S2.** Symbols used in this article.

| Reaction | Reference |
|---|---|
| ***1. Calvin Cycle*** | |
| 2 $CO_2$ + 7 ATP + 4 NADH → 1 Acetyl-CoA | Salimijazi *et al*. [Salimijazi2020b]. |
| 3 $CO_2$ + 7 ATP + 5 NADH → 1 Pyruvate | Salimijazi *et al* [Salimijazi2020b]. |
| ***2. Wood-Ljungdahl Pathway*** | |
| 4 $CO_2$ + 2 ATP + 8 NADH → 2 Acetyl-CoA | Berg [Berg2011a]. |
| 2 $Fd_{red}$ + Acetyl-CoA + $CO_2$ → Pyruvate | KEGG R01196. |
| ***3. Reductive TCA Cycle*** | |
| 4 CO2 + 4 ATP + 8 NADH → 2 Acetyl-CoA | Alissandratos *et al.* [Alissandratos2015a], Claassens *et al*. [Claassens2016a]. |
| 2 $Fd_{red}$ + Acetyl-CoA + $CO_2$ → Pyruvate | KEGG R01196. |
| ***4. 3-hydroxypropionate/4-hydroxybutyrate Cycle*** | |
| 6 $HCO_3^-$ + 10 ATP + 10 NADH → 2 pyruvate | Berg *et al*. [BergI2007a], Claassens *et al*. [Claassens2016a]. |
| 2 Pyruvate → 2 Acetyl-CoA + 2 NADH + 2 $CO_2$ | Berg [Berg2002a], Schomburg *et al.* [Schomburg2017a]. |
| ***5. 3-hydroxypropionate Cycle*** | |
| 6 $HCO_3^-$ + 10 ATP + 12 NADH → 2 Pyruvate | Zarzycki *et al*. [Zarzycki2009a], Herter *et al*. [Herter2002a], Berg [Berg2002a]. |
| 2 Pyruvate → 2 Acetyl-CoA + 2 NADH + 2 $CO_2$ | Zarzycki *et al*. [Zarzycki2009a], Herter *et al*. [Herter2002a], Berg [Berg2002a]. |
| ***6. 4-hydroxybutyrate Cycle*** | |
| 1 $CO_2$ + 1 $HCO_3^-$ + 3 ATP + 1 NADH + 6 $Fd_{red}$ → 1 Acetyl-CoA | Huber *et al*. [Huber2008a]. |
| 2 Pyruvate → 2 Acetyl-CoA + 2 NADH + 2 $CO_2$ | Berg [Berg2002a], Schomburg *et al.* [Schomburg2017a]. |
| ***7. Formolase Pathway*** | |
| 6 $HCO_2^-$ + 10 ATP + 4 NADH → 2 3-PG | Siegel *et al*. [Siegel2015a], Bar-Even *et al*. [Bar-Even2016a]. |
| 2 3-PG → 2 Pyruvate + 2 ATP | Berg [Berg2002a]. |
| 2 Pyruvate → 2 Acetyl-CoA + 2 NADH + 2 $CO_2$ | Berg [Berg2002a], Schomburg *et al.* [Schomburg2017a]. |

**Table S3.** $CO_2$-fixation and $C_1$-assimilation reactions. $CO_2$-fixation and $C_1$-assimilation reactions considered in this article were first assembled in Salimijazi *et al*. [Salimijazi2020b] and are restated here for convenience. Overall reactions for production of metabolic intermediates by 6 naturally-occurring $CO_2$-fixation cycles and the synthetic Formolase formate assimilation pathway, and the FeMoCo nitrogenase $N_2$-fixation reaction. Reactions can be referenced KEGG database [Kanehisa2000a, Kanehisa2019a, Kanehisa2021a]. $Fd_{red}$: Reduced Ferredoxin; 3-PG: 3-Phosphoglycerate.

| # | Reaction | EC | KEGG |
|---|---|---|---|
| 1 | Acetyl-CoA + Pyruvate + $H_2O$ → (R)-2-Methylmalate + CoA | 2.3.3.21 | R07399 |
| 2 | (R)-2-Methylmalate → 2-Methylmaleate + $H_2O$ | 4.2.1.35 | R03896 |
| 3 | 2-Methylmaleate + $H_2O$ → D-erythro-3-Methylmalate | 4.2.1.35 | R03898 |
| 4 | D-erythro-3-Methylmalate + NAD+ → 2-Oxobutanoate + $CO_2$ + NADH + $H^+$ | 1.1.1.85 | R00994 |
| 5 | 2-Oxobutanoate + CoA → Propionyl-CoA + Formate | 2.3.1.54 | R06987 |
| 6 | Formate → $CO_2$ + NADH + $H^+$ | 1.17.1.9 | R00519 |

**Table S4.** Acetyl-CoA to propionyl-CoA reactions.

### Supplementary Information References


[Alissandratos2015a] A. Alissandratos and C. J. Easton. “Biocatalysis for the application of $CO_2$ as a chemical feedstock”. *Beilstein Journal of Organic Chemistry* 11 (2015), pp. 2370–2387. doi:10.3762/bjoc.11.259.

[Bar-Even2016a] A. Bar-Even. “Formate Assimilation: The Metabolic Architecture of Natural and Synthetic Pathways.” *Biochemistry* 55 (2016), pp. 3851–63. doi:10.1021/acs.biochem.6b00495.

[Berg2002a] J. Berg, J. Tymoczko, and L. Stryer. Biochemistry. 5th. New York, NY: W H Freeman, 2002.

[Berg2011a] I. A. Berg. “Ecological Aspects of the Distribution of Different Autotrophic $CO_2$ Fixation Pathways”. *Applied and Environmental Microbiology* 77 (2011), pp. 1925–1936. doi:10.1128/aem.02473-10.

[BergI2007a] I. A. Berg, D. Kockelkorn, W. Buckel, and G. Fuchs. “A 3-Hydroxypropionate/4-Hydroxybutyrate Autotrophic Carbon Dioxide Assimilation Pathway in Archaea”. *Science* 318 (2007), pp. 1782–1786. doi:10.1126/science.1149976.

[Claassens2016a] N. J. Claassens, D. Z. Sousa, V. A. M. dos Santos, W. M. de Vos, and J. van der Oost. “Harnessing the power of microbial autotrophy”. *Nature Reviews Microbiology* 14 (2016), pp. 692–706. doi:10.1038/nrmicro.2016.130.

[Herter2002a] S. Herter, G. Fuchs, A. Bacher, and W. Eisenreich. “A bicyclic autotrophic $CO_2$ fixation pathway in *Chloroflexus aurantiacus*”. *Journal of Biological Chemistry* 277 (2002), pp. 20277–20283. doi:10.1074/jbc.m201030200.

[Huber2008a] H. Huber, M. Gallenberger, U. Jahn, E. Eylert, I. A. Berg, D. Kockelkorn, W. Eisenreich, and G. Fuchs. “A dicarboxylate/4-hydroxybutyrate autotrophic carbon assimilation cycle in the hyperthermophilic *Archaeum Ignicoccus hospitalis*”. *Proceedings of the National Academy of Sciences* 105 (2008), pp. 7851–7856. doi:10.1073/pnas.0801043105.

[Kanehisa2000a] M. Kanehisa and S. Goto. “KEGG: Kyoto Encyclopedia of Genes and Genomes”. *Nucleic Acids Research* 28.1 (2000), pp. 27–30. doi:10.1093/nar/28.1.27.

[Kanehisa2019a] M. Kanehisa. “Toward understanding the origin and evolution of cellular organisms”. Protein *Science* 28.11 (2019), pp. 1947–1951. doi:10.1002/pro.3715.

[Kanehisa2021a] M. Kanehisa, M. Furumichi, Y. Sato, M. Ishiguro-Watanabe,and M. Tanabe. “KEGG: integrating viruses and cellular organisms”. *Nucleic Acids Research* 49.D1 (2020), gkaa970–. doi:10.1093/nar/gkaa970.

[NIST2022a] P.J. Linstrom and W.G. Mallard, Eds., NIST Chemistry WebBook, NIST Standard Reference Database Number 69, National Institute of Standards and Technology, Gaithersburg MD, 20899 (retrieved September 24, 2022). doi:10.18434/T4D303.

[Salimijazi2020b] F. Salimijazi, J. Kim, A. M. Schmitz, R. Grenville, A. Bocarsly, and B. Barstow. “Constraints on the Efficiency of Engineered Electromicrobial Production”. *Joule* 4 (2020), pp. 2101–2130. doi:10.1016/j.joule.2020.08.010.

[Schomburg2017a] I. Schomburg, L. Jeske, M. Ulbrich, S. Placzek, A. Chang,and D. Schomburg. “The BRENDA enzyme information system–From a database to an expert system”. *Journal of Biotechnology* 261 (2017), pp. 194–206. doi:10.1016/j.jbiotec.2017.04.020.

[Siegel2015a] J. B. Siegel, A. L. Smith, S. Poust, A. J. Wargacki, A.Bar-Even, C. Louw, B. W. Shen, C. B. Eiben, H. M.Tran, E. Noor, J. L. Gallaher, J. Bale, Y. Yoshikuni, M. H. Gelb, J. D. Keasling, B. L. Stoddard, M. E. Lidstrom, and D. Baker. “Computational protein design enables a novel one-carbon assimilation pathway”. *Proceedings of the National Academy of Sciences* 112 (2015), p. 3704-3709. doi:10.1073/pnas.1500545112.

[Zarzycki2009a] J. Zarzycki, V. Brecht, M. Müller, and G. Fuchs. "Identifying the missing steps of the autotrophic 3-hydroxypropionate $CO_2$ fixation cycle in *Chloroflexus aurantiacus*". *Proceedings of the National Academy of Sciences* 106 (2009), p. 21317. doi.10.1073/pnas.0908356106.
